# Transposon expression in the *Drosophila* brain is driven by neighboring genes and diversifies the neural transcriptome

**DOI:** 10.1101/838045

**Authors:** Christoph D. Treiber, Scott Waddell

**Affiliations:** Centre for Neural Circuits and Behaviour, University of Oxford, Tinsley Building, Mansfield Road, Oxford OX1 3SR, UK

**Keywords:** Transposon expression, alternative splicing, transcriptional heterogeneity, single-cell transcriptomics.

## Abstract

Somatic transposition in neural tissue could contribute to neuropathology and individuality, but its prevalence is debated. We used single-cell mRNA sequencing to map transposon expression in the *Drosophila* midbrain. We found that neural transposon expression is driven by cellular genes. Every expressed transposon is resident in at least one cellular gene with a matching expression pattern. A new long-read RNA sequencing approach revealed that coexpression is a physical link in the form of abundant chimeric transposon-gene mRNAs. We identified 148 genes where transposons introduce cryptic splice sites into the nascent transcript and thereby produce many additional mRNAs. Some genes exclusively produce chimeric mRNAs with transposon sequence and on average transposon-gene chimeras account for 20% of the mRNAs produced from a given gene. Transposons therefore significantly expand the neural transcriptome. We propose that chimeric mRNAs produced by splicing into polymorphic transposons may contribute to functional differences between individual cells and animals.

## Introduction

Transposons comprise almost half of every eukaryote genome (Britten and Kohne, 1968; International Human Genome Sequencing Consortium et al., 2001; Ketchum et al., 2000) and their mobilization in the germline contributes to chromosome evolution. Non-heritable *de novo* transposon activity in neural tissue has been proposed to contribute to functional heterogeneity in the brain and to neurological disease (Baillie et al., 2011; Coufal et al., 2009; Evrony et al., 2012; Kazazian, 2011; Kazazian and Moran, 2017; Muotri et al., 2005; Schauer et al., 2018). However, it is difficult to faithfully map rare *de novo* transposon insertions using whole-genome DNA sequencing (Baillie et al., 2011; Evrony et al., 2012, 2016; Perrat et al., 2013; Treiber and Waddell, 2017; Upton et al., 2015). A growing number of studies have therefore correlated the development of neurodegeneration in animal models with changes in transposon expression (Guo et al., 2018; Krug et al., 2017; Li et al., 2013; Li et al., 2012; Sun et al., 2018). Using expression as a proxy for mobility could be misleading because high-level transposon expression does not appear to result in elevated *de-novo* somatic transposition in the brain (Evrony et al., 2012, 2016; Treiber and Waddell, 2017). It is therefore important to understand what controls the expression of transposon-derived sequences in the brain and whether their elevated expression relates to neural function.

An early study of the human LINE-1 (L1) promoter demonstrated that its activity was heavily influenced by flanking cellular sequences, and concluded that expression of a given L1 depended on its location in the genome (Lavie et al., 2004). Such a locus specific model could more generally explain the apparent cell-type restricted nature of transposon expression and mobilization in the brain. So far, studies of somatic transposon expression have either focused on single transposon families or have been based on bulk sequencing of tissues or cultured cells (Chung et al., 2019; Faulkner et al., 2009; Li et al., 2013; Philippe et al., 2016; Rangwala et al., 2009). However, answering this question on a brain- and genome-wide scale requires a means to relate the cellular expression of each transposon in the genome to that of their neighboring genes. Recent technical developments in whole-genome DNA sequencing and high-throughput single-cell transcriptomics of complex tissues now make this possible (Macosko et al., 2015).

Here we used single-cell transcriptomics to map transposon expression to individual cells in the *Drosophila* midbrain. We found that many transposons are expressed with cell-specificity that is often highly correlated to that of a neighboring gene. A more detailed analysis revealed that >90% of transposon expression can be linked to co-expression with a host gene, indicating that these genes are the main driver of somatic transposon expression. Long-read sequencing showed that the transposon and neighboring gene are alternatively spliced together becoming part of the same chimeric mRNAs. Sometimes all mRNAs produced from a particular gene in which a transposon resides include transposon sequence. Therefore, transposons produce genome-wide diversification of cellular transcripts. Analysis of sequencing data produced from other fly strains demonstrated large differences in their chimeric transcriptomes. Inter-strain and individual differences in transposon complement therefore constrain the cellular specificity and likelihood of transposon-directed pathology.

## Results

### Single-cell transcriptomics reveals cell-type restricted transposon expression

The *Drosophila* genome contains 112 families of transposons and the number of an individual type varies from a few to hundreds of copies (Kaminker et al., 2002). Conventional single-cell RNAseq (scRNAseq) analysis pipelines typically discard sequencing reads that align to multiple genomic loci, and so they overlook transposon expression. We therefore devised an analysis pipeline to map the expression of all transposons within scRNAseq data. We masked all repetitive sequences in the reference genome and then added a single copy of the consensus sequence for every known transposon to the masked genome. In essence this produces a *Drosophila* reference genome with one copy of each type of transposon. We first used this modified reference genome to map transposon expression onto single cells of the midbrain prepared from a fly strain expressing mCherry in αβ Kenyon cells (KCs) of the mushroom body (MB); from here called αβCherry flies. We found evidence for expression of both the sense and the antisense strand of most transposons, which comprised 76.2 and 23.8% (+/− 1.7% SD) of all transposon expression, respectively (Supplemental figure 1). We first performed principal component decomposition of cellular genes and clustered cells from the midbrain by constructing a k-Nearest-Neighbor graph on the Euclidean distances in the PCA space, optimizing the modularity using the Louvain algorithm (Butler et al., 2018). This analysis grouped cells into many discrete clusters. We next assigned many of these clusters to cell types in the midbrain using the expression patterns of known marker genes (Croset et al., 2018) (Figure 1a). Displaying the expression of individual types of transposons on the cluster plot revealed that some transposons are up-regulated in specific cell types. For example, the long-terminal repeat (LTR) retrotransposons COPIA and NOMAD showed elevated expression in the αβ, *α*′*β*′ and *γ* Kenyon Cells (KCs) classes (Figure 1b, first graph) and *α*′*β*′ KCs (Figure 1c, first graph), respectively. Other LTR retrotransposons such as MICROPIA were upregulated in the ellipsoid body (Figure 1d, first graph) whereas BLOOD and 412 were higher in glia (Figure 1e,f, first graphs).

**Figure 1.**
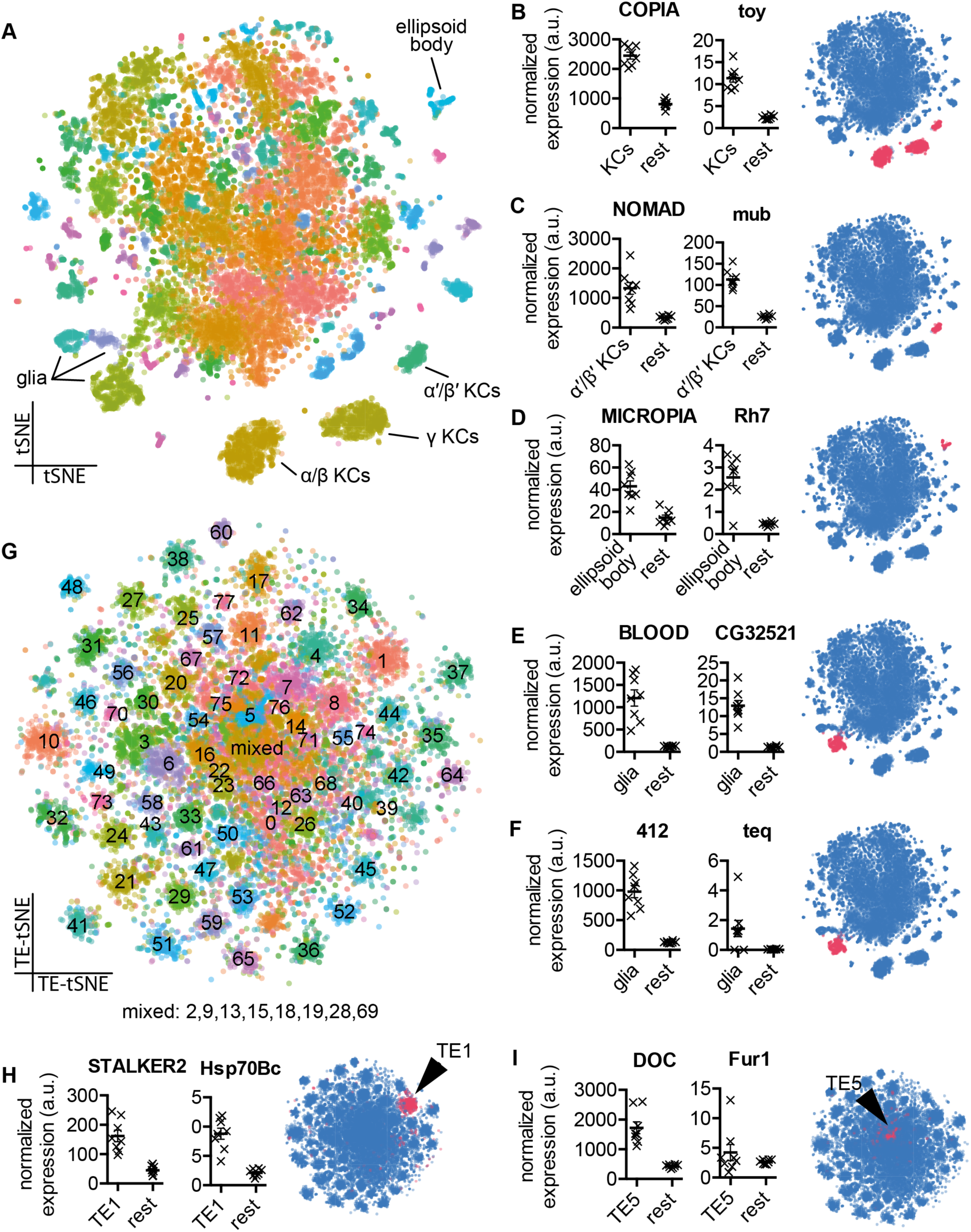
Single-cell transcriptomics reveals patterned transposon expression in the *Drosophila* midbrain. **A** Two-dimensional reduction (tSNE) of 14,804 *Drosophila* midbrain cells, based on gene expression levels. Colors represent cell clusters (at SNN resolution of 3.5). **B-F** Mean expression of transposons and neighboring cellular genes in the relevant cell groups in 8 biological replicates and tSNE representation of cell-type restricted expression. **B** COPIA and *twin-of-eyeless* (*toy*) in all Kenyon Cell (KC) classes. **C** NOMAD and *mushroom-body expressed* (*mub*) in *α*′*β*′ KCs. **D** MICROPIA and *Rhodopsin 7* (*Rh7*) in the ellipsoid body **E and F** BLOOD and *CG32521*, and 412 and *tequila* (*teq*) in glia. Values represent the mean normalized number of unique molecular identifiers (UMI’s) in an average cell from each cell type, and from the rest of the midbrain. Error bar indicates standard error of mean (SEM). Note that transposon- and gene levels were normalized separately. Blue schematic shows location of cell cluster (pink) in tSNE plot. **G** Two-dimensional reduction of 14,804 *Drosophila* midbrain cells, based exclusively on transposon expression levels. Colors represent cell clusters (at SNN resolution of 3.5). **H and I** Mean expression of STALKER2 and *Hsp70Bc* and DOC and *Furin 1* (*Fur1*) in their relevant transposon clusters and the position of the cluster in the overall transposon-based tSNE (indicated in pink).

### Transposon expression correlates with that of cellular genes they are inserted within

Transposons could be elevated in specific cell types because they are inserted in genes that are highly expressed in the same cells. To test this hypothesis, we re-used our previously published high-coverage gDNA sequence of αβCherry flies. We mapped all the germline transposon insertions in these flies using TEchim, a new custom-built transposon analysis program. TEchim first generates long nucleotide contigs from either gDNA or cDNA sequencing reads, then creates in-silico paired-end reads and screens them for cases where one in-silico end maps to a cellular gene and the mate read maps to a specific transposon. Since these paired-end reads are derived from contiguous sequences, TEchim permits one to subsequently refer back to the original long reads to determine the precise nucleotide sequence of the transposon-gene breakpoints. Using TEchim, we found copies of COPIA, NOMAD, MICROPIA, BLOOD and 412 inside the genes *twin-of-eyeless* (*toy*), *mushroom-body expressed* (*mub*), *Rhodopsin-7* (*Rh7*), *CG32521* and *tequila* (*teq*), respectively. The expression of each of these genes mirrored the expression pattern of the transposon they harbored (Figure 1b-f, second graphs). The expression of these transposons in the brain of αβCherry flies therefore appears to be driven by their relevant host genes.

We next tested whether all transposons exhibit patterned expression throughout the midbrain. We re-clustered the single-cell data of the fly using only transposon expression. This generated 78 cell clusters that mostly contained cells from all 8 biological replicates in the data (Figure 1g, Supplemental figure 2). This result suggests that transposon expression is stereotyped across samples derived from different flies collected from the same strain. We then analyzed the expression of cellular genes across the transposon clusters and found many clusters also preferentially expressed certain genes. For example, the cluster of cells expressing the LTR of STALKER2 was enriched for cells that also expressed the *Hsp70Bc* gene (Figure 1h), and cells in the DOC-positive cluster showed increased expression of *Furin1* (*Fur1*). By referring back to the gDNA, we found that αβCherry flies harbor a copy of STALKER2 within *Hsp70Bc* and a copy of the LINE-like DOC element inside *Fur1*. Again, these data suggest that expression of STALKER2 and DOC is driven by a neighbouring gene.

### Quantitative analysis reveals high fidelity transposon-gene co-expression

Our gDNA analysis also revealed many transposon insertions inside genes that were more broadly expressed across the brain. In total, we identified 1952 germline transposon insertions within genes. Of these, 881 cases were inserted in the sense direction to the open reading frame of the gene and 1071 were in the antisense orientation (Supplemental Table 1). To quantify the correlated expression of transposons and cellular genes we devised a method based on the established Hardy-Weinberg principle for quantifying linkage equilibrium of two alleles in population genetics (Lewontin and Kojima, 1960) (Figure 2a). We first binarized our scRNAseq data to generate the equivalent of bi-allelic traits in a population. We then calculated the proportion of cells expressing a specific transposon, multiplied it by the proportion of cells expressing a certain gene, and then subtracted this value from the proportion of cells that expressed both the transposon and the gene. We termed this value the Coexpression Disequilibrium, CD. We also normalized these CD values to account for the variable abundance of each transposon and gene in every transposon-gene pair and repeated the analysis for all transposon-transposon and gene-gene pairs. These normalized values were then ranked within each of the 8 biological replicates. P-values were corrected for multiple comparisons and describe the probability that a transposon-gene pair would have such a highly ranked CD value across multiple replicates if they were expressed independently.

**Figure 2.**
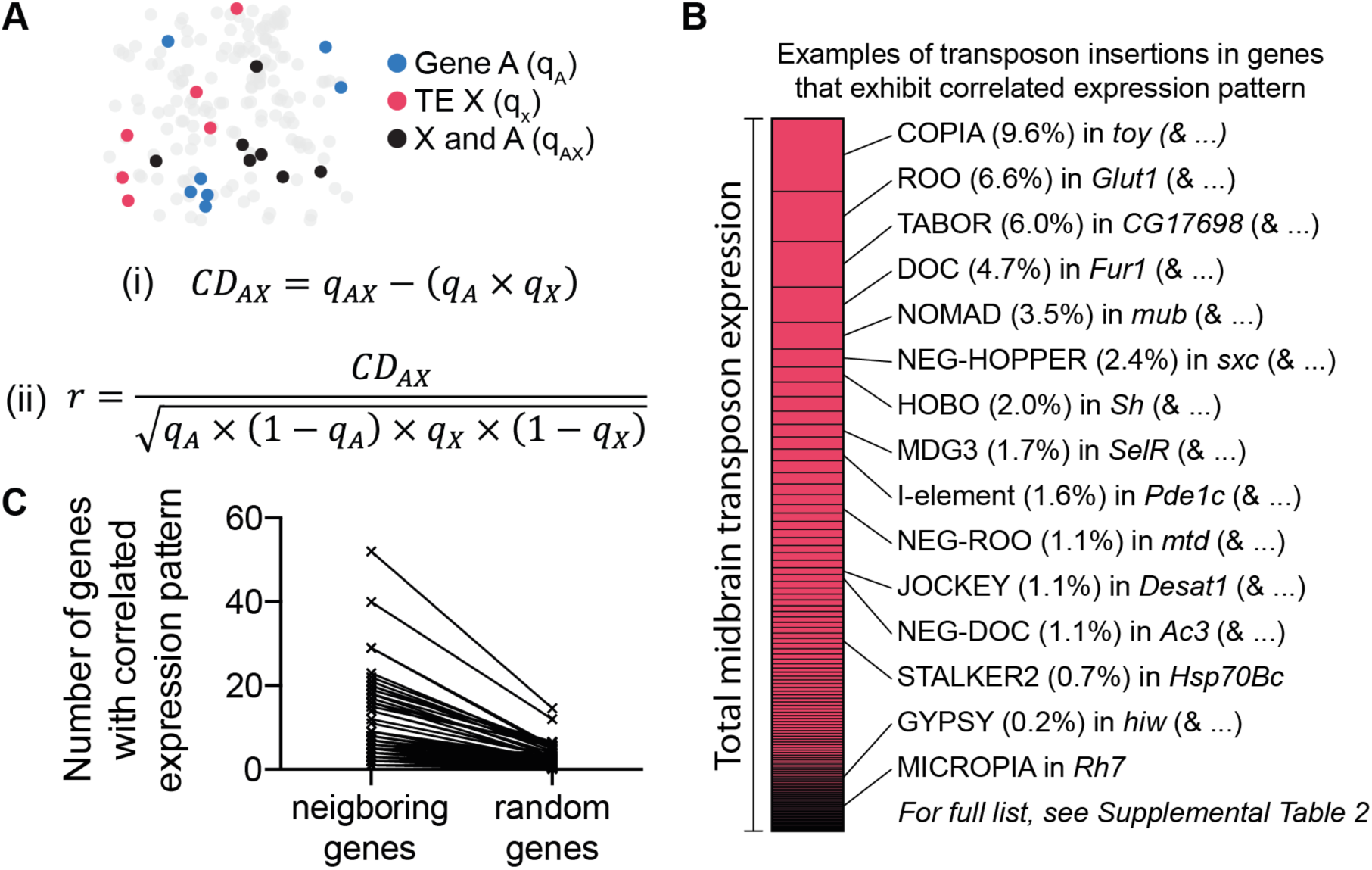
Transposons are co-expressed with neighboring genes. **A** Schematic and formulae describing the calculation of Co-expression Disequilibrium values. **B** Examples of transposon-gene pairs that are neighboring in the genome and co-expressed across the midbrain. Pink bar represents the total transposon expression. Examples are selected from the most highly expressing transposons. See Supplemental Table 2 for entire list of correlated transposon-gene pairs. **C** Graph illustrating that more neighboring genes are correlated with their resident transposons than if transposons are randomly assigned to genes.

We combined the list of all germline transposon insertions in αβCherry flies with the scRNAseq data generated from the same population of flies and calculated the CD values between every transposon and the gene in which it was inserted. We tested every transposon that contributed at least 0.1% of the overall transposon expression in our midbrain αβCherry fly samples. This cut-off left 59 different transposons, 34 of which were expressed in both the sense- and the antisense direction, 22 only in sense, and 3 only in antisense (Figure 2b, Supplemental Table 2). For 56 of these transposons we found at least one copy inside a gene that exhibited a correlated expression pattern (Benjamini-Hochberg corrected p-val >0.05). For those cases where the transposon was inserted in the same orientation as the transcription unit of the gene the expression of the sense strand of the transposon correlated to that of the gene. In contrast, the antisense strand of reverse orientation transposons was correlated with the host gene. Importantly, the average number of correlated genes for each transposon was significantly less if equivalent values were calculated using random assignment of transposons to host genes (Figure 2c). These analyses therefore demonstrate that the genomic locus strongly influences the expression patterns of almost all transposons in the fly brain. We did not identify a neighboring gene with a correlated expression pattern for the transposons TART-A, P-element and HMSBEAGLE. This is expected for TART-A, which is a telomeric retrotransposon, and for P-element, which is a remnant of transgenic intervention in αβCherry flies. It is conceivable that the HMSBEAGLE retrotransposon is the only element that is expressed independently of a cellular gene. However, despite our high coverage sequence of αβCherry flies we may have missed a germline HMSBEAGLE insertion that sits near a gene that is driving its expression.

### Transposons are exonized into cellular mRNAs

Recent work has shown that mRNAs from the *Arc* gene contain transposon-like sequence in the coding sequence and 3′ UTR (Pastuzyn et al., 2018; Zhang et al., 2015). We therefore tested whether chimeric mRNAs might occur more broadly and extend to all transposons. We extracted mRNA from αβCherry fly heads and generated 250 basepair long reads which were screened using TEchim for chimeric reads. We also incorporated a function in TEchim that maintains strand-specificity of the input reads which enabled us to unambiguously assign chimera to cellular genes. This analysis revealed that a large number of transposons inside introns lead to the formation of chimeric mRNAs. In total, we found chimeric mRNA from 887 transposon insertions (Figure 3a, Supplemental Table 3). Chimera included sequences from LTR, LINE-like and DNA transposons attached to mRNAs from genes involved in a broad range of biological processes. For example, we found sequence from the LTR-retrotransposon GYPSY in transcripts of the ubiquitin gene *Ubi-P5E* and of the neuron specific ubiquitin ligase *highwire* (*hiw*), the non-LTR element DOC in *Fur1*, encoding a synaptic membrane bound protease, and the TIR element HOBO attached to transcripts from *Shaker*, which encodes a voltage-gated potassium channel (Izquierdo, 1994; Kaplan and Trout, 1969; Roebroek et al., 1991; Wan et al., 2000).

**Figure 3.**
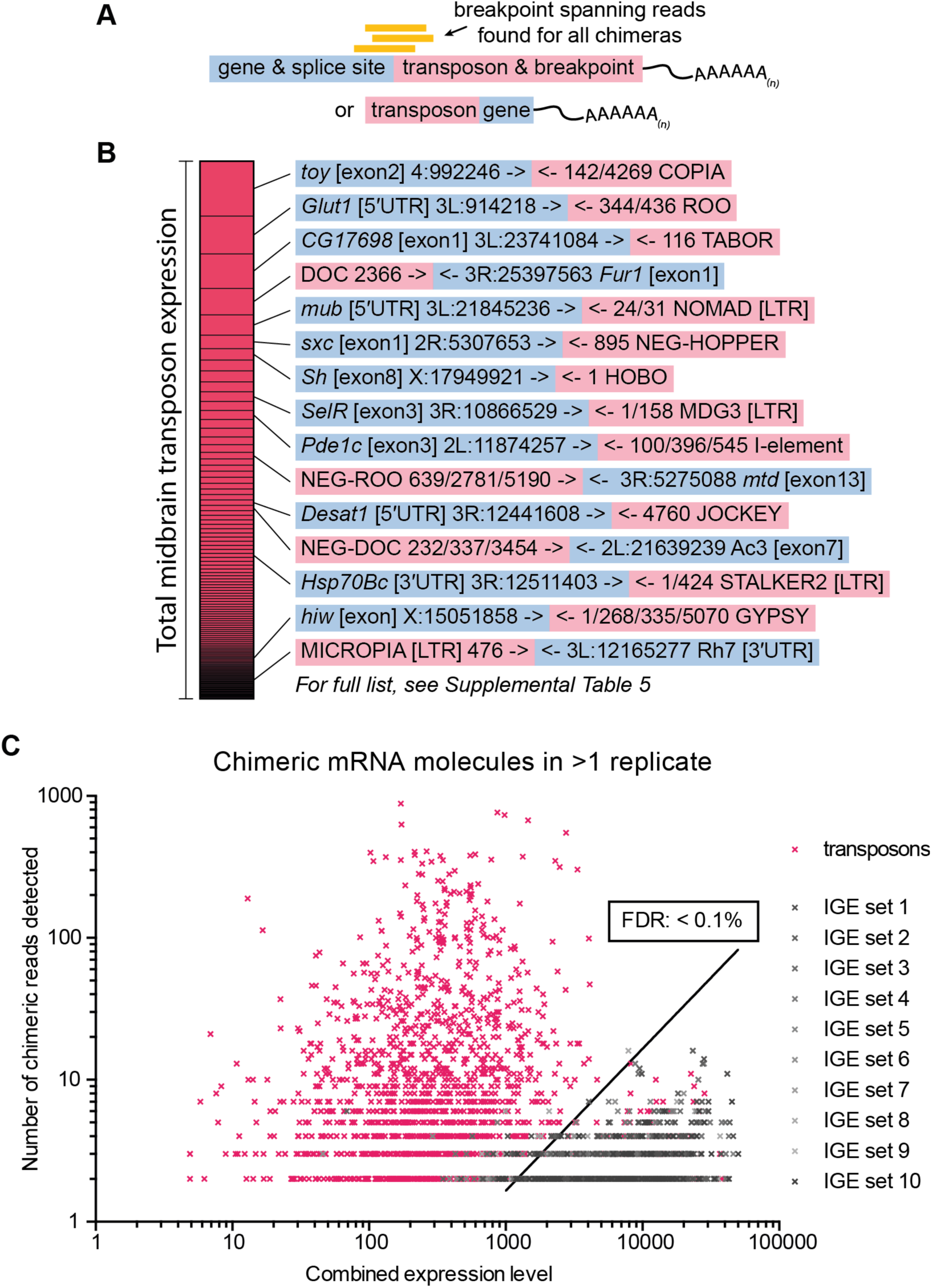
Chimeric transposon-gene mRNA is abundant in the midbrain. **A** Schematic of the structure of chimeric mRNA molecules, and illustrating the data representation in **B**. **B** Examples of transposon-gene pairs that form chimeric mRNA. The blue box contains the gene name, the section of the gene that forms a chimera and the breakpoint, which is always the endogenous exon-intron junction. The red box contains the transposon name, and the breakpoint(s) on the transposon. Note that for consistency, transposon breakpoints are always taken from the sense orientation, starting from the core transposon sequence, unless specifically indicated as [LTR]. Examples shown here are the same transposon-gene pairs as in Figure 2b. For the entire list of chimera, see Supplemental Table 5. **C** Graph showing the number of chimeric reads, and the combined expression levels of each transposon-gene pair (pink), as well as for all 10 sets of IGE-gene pairs (grey). Combined expression levels are the square root of the product of reads in both transcripts of a transposon/IGE-gene pair. IGEs were used to calculate a threshold for a False Discovery Rate that is less than 0.1%.

Previous studies of chimeric sequencing reads have established that *in vitro* amplification of genetic material often leads to chimeric amplification artefacts (Evrony et al., 2016; Treiber and Waddell, 2017). It was thus important to account for similar errors in our data. We therefore calculated the rate of these artefacts in our mRNA data by selecting 10 sets of 167 exons (with each set providing a number of sequences corresponding to the 112 different transposon types and 55 LTRs) with matching expression levels in the brain. These exons lack the ability to relocate in gDNA so we refer to them as immobile genetic elements (IGEs). Since IGEs should only occur as single copies in the gDNA from αβCherry, chimeric reads between IGEs and other genes most likely represent amplification artefacts. We found the rate of generating IGE chimeras was directly correlated to the expression level of the IGE and the gene that it formed a chimeric molecule with. Critically, the IGE chimera rate was substantially lower than that of chimera formed between genes and transposons. We therefore used the rate of IGE chimera to filter the transposon chimera detected using TEchim, defining a false discovery rate (FDR) of 0.1% (Figure 3b, Supplemental Table 3). All examples of chimeric transcripts that are presented in detail in this study have supporting evidence that exceeds this FDR.

### LTR retrotransposon expression is predominantly non-autonomously

Given that transposon expression was highly correlated with at least one neighboring gene, we hypothesized that all transposon expression in the brain might occur as co-expression with cellular genes. To test this, we focused on LTR retrotransposons. We quantified the number of reads that spanned an LTR-gene breakpoint, and compared it to the number of reads that crossed the LTR-transposon breakpoint within the transposon (Figure 4a). Autonomously expressed, full-length transposons only generate the latter type of read, whilst non-autonomous expression should generate both kinds of reads, at varying proportions. We found that around 95% of the highly expressed LTR transposons produced a roughly equivalent number of gene-transposon and LTR-transposon reads, suggesting that LTR transposons are expressed as chimeras with cellular genes, rather than being autonomously expressed (Figure 4b, Supplemental Table 4).

**Figure 4.**
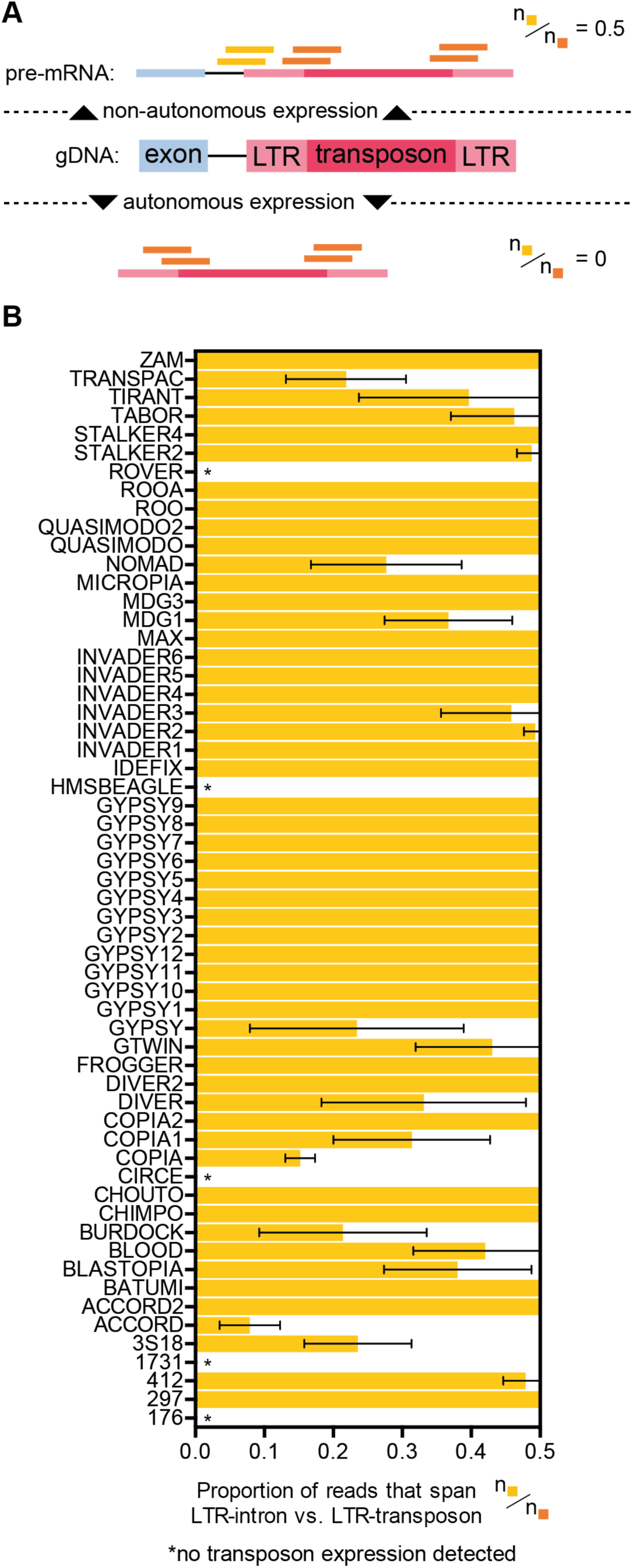
LTR retrotransposon expression is predominantly non-autonomous. **A** Illustration showing method of calculating the percentage of chimeric transcripts vs. autonomously expressed transposon transcripts. Non-autonomous expression should result in an approximate value of 0.5 for reads spanning the LTR section of transposons and the intron of a neighboring gene over the number of reads spanning the LTR and the core section of the transposon. In contrast, autonomous expression would not result in LTR-gene spanning reads. **B** List of all LTR transposons analyzed in our mRNA data. We identified LTR-gene spanning reads for every LTR transposon that is expressed in the midbrain. Error bars represent standard deviation. Values have been capped at 0.5 in this graph. However, some transposons produced a much higher number of LTR-gene reads (see Supplemental Table 4).

### Many transposons introduce cryptic alternative splice-sites into cellular genes

Given the abundance of reads that span a gene-transposon breakpoint, we next investigated the structure of these transcripts in more detail. Our transposon mapping identified 887 loci with a germline transposon insertion in the intron of a gene. For each of these we found mRNA molecules where one section mapped to the beginning or end of the transposon and the other section corresponded to the flanking intronic sequence. These reads could represent nascent unspliced chimeric pre-mRNAs from which transposon-derived sequence could be removed by splicing to yield intact host mRNAs, and full-length transposon sequences. However, we also found 148 examples where breakpoint-spanning reads indicate that specific sections of transposon sequence are spliced into host-gene transcripts (Supplemental Table 5).

Analysis of the breakpoints inside transposons at these 148 sites revealed that chimera are formed at conserved locations in each type of transposon. For example, in cases where an antisense ROO resided within an intron, we found transcripts where the 3′-end of an upstream exon had formed a new phosphate bond to a section of ROO at positions 5460 and 2094 at 19 and 3 different genomic loci, respectively, and also at several additional breakpoints with lower frequency (Figure 5a,b). In addition, we identified transcripts where sections of ROO were bound to the 5′-end of a downstream exon. We found breakpoints at position 5191 from 24 genes, two at 2783, and several others at unique positions (note that the numbering runs backward because it relates to the forward orientation of ROO). Whereas intronic antisense ROO provides gene-transposon breakpoints for 28 exons, and transposon-gene breakpoints for 33, intronic sense ROO only introduced 4 and 1 (Figure 5c). Similarly, the LTR BLOOD also introduced more breakpoints when it was inserted in the antisense orientation relative to the host gene (14 vs. 6, Figure 5d,e).

**Figure 5.**
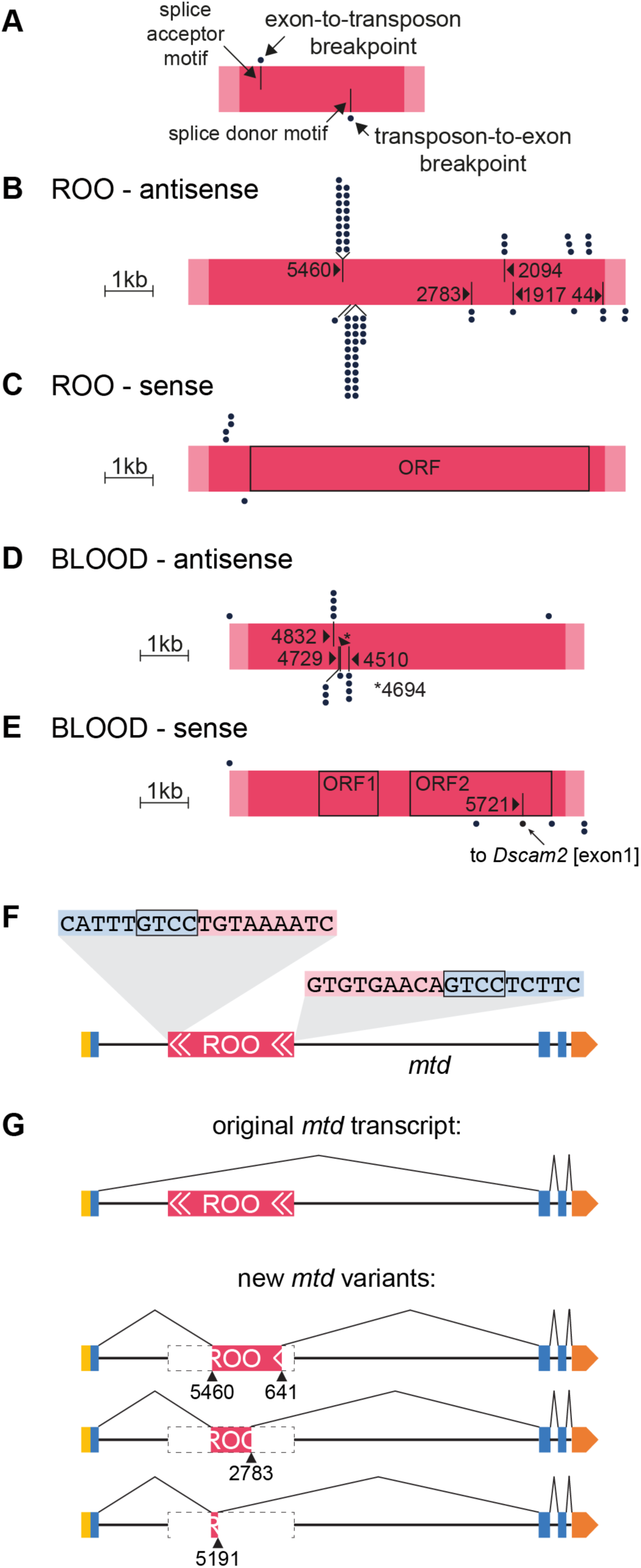
Transposons introduce splice sites at conserved locations. **A** Illustration of the labelling scheme in panels **B-E**. The pink bar represents the transposon; light pink ends indicate the LTRs and the dark pink the core sequence. The positions of the dots above the bar represent the site on the transposon where an upstream exon splice donor site has merged. Every dot represents a different gene. Black lines in the top half of the pink bar represent splice acceptor (SA) motifs in the transposon. Dots below the pink bar indicate the location of breakpoints on the transposon that are spliced to upstream exonic SA sites of different genes. Bars in the lower half indicate splice donor (SD) motifs. **B-E** Representations of sense- and antisense ROO and BLOOD (to scale), with all breakpoints to SA and SD sites of neighboring genes. Note that the frequently used site on antisense ROO at position 5191 is a non-consensus SD site, which lacks the expected GT motif at the immediate breakpoint. The sequence around 5191 resembles the consensus SD motif, although the GT is a GC. Compare TTTGGCAAGTT to motif in Supplemental figure 3a. **F** Illustration of antisense ROO insertion in the *mustard* (*mtd*) gene. Only one isoform of *mtd* is shown. Yellow box represents the 5′UTR, blue boxes are exons, orange box the 3′UTR, pink represents ROO transposon with white arrows indicating the LTRs. Breakpoint-spanning gDNA reads reveal Target Site Duplication (TSD, inset). **G** Schematic of original *mtd* transcript, and of three new splice isoforms.

We screened the transposon sections around breakpoints for consensus splice-acceptor (SA) and donor (SD) sequence motifs and found many cases where the gene-to-transposon chimera had been formed at SA consensus motifs, and transposon-to-gene chimera at SD motifs (Stephens and Schneider, 1992) (Supplemental figure 3, Supplemental Table 6). For example, all breakpoints in antisense BLOOD that were formed with more than one exon were precisely located at the predicted SA and SD splice site (Figure 5d). Interestingly, we did not find a consensus SD motif at the transposon-gene breakpoint at position 5191 of antisense ROO, although it frequently provided 5′-sequence to transposon-gene chimeric RNAs. However, the sequence around position 5191 matched the consensus motif, with the exception of a GT-to-GC conversion (see Supplemental figure 3). Taken together, our analysis revealed that transposons introduce many alternative splice sites, which are recognized by the host cell spliceosome to combine cellular exonic sequences with sections of transposon sequence.

We also identified cases of alternative splicing to different sites within the same transposon insertion. Again using ROO as an example, αβCherry flies harbor a reverse orientation ROO in the intron between exons 10 and 11 of the pan-neurally expressed *mustard* (*mtd*) gene, which to date has only been implicated in fly innate immunity (Wang et al., 2012) (Figure 5f). RNAseq revealed a complex collection of *mtd* splice variants that incorporated different fragments of ROO (Figure 5g). SD sites upstream of this ROO came from the end of either *mtd* exon 11 or 13. and these spliced in to the corresponding SA at position 5462 within ROO (Figure 5g). We also found three different SD sites (at positions 641, 2784 and 5191) within ROO, which spliced out to the closest downstream SA (exon 6) of *mtd*. Therefore, this ROO element substantially increases the *mtd* mRNA isoform repertoire. Without ROO the *mtd* locus can express 23 isoforms whereas with ROO it can generate 68 differentially spliced mRNAs.

We identified 147 other genes whose transcript diversity was similarly increased by transposon insertions. These alternative transcripts incorporate 43 different transposon families which each introduce cryptic SA and/or SD sites into host genes (see Supplemental Table 5). For example, we found a sense insertion of BLOOD inside the *Dscam2* gene which encodes the transmembrane Down Syndrome cell adhesion molecule 2. Chimeric reads indicate that transcription of *Dscam2* is frequently initiated in BLOOD, which is then spliced into exon 33 (the second exon) of the gene. This splicing event combines ORF2 of BLOOD with the remaining exons of *Dscam2* and correctly aligns the reading frames of the two transcripts, generating a novel N-terminus (Supplemental figure 4). We also observed cases where transposon chimera resulted in exon skipping (e.g. the above mentioned ROO in *mtd* and 412 inside *tequila*, Supplemental figure 5). Most transposon chimera resulted from intronic insertions, but we also detected one case where a HOBO insertion resided in the exon of the *CG31705* gene. This HOBO introduced a cryptic SA site which was spliced to the upstream SD from the first exon, creating a truncated *CG31705* transcript (Supplemental figure 6). Together these data show that a broad range of *Drosophila* transposons are alternatively spliced into mRNAs producing many more isoforms of a large number of neurally expressed genes.

### Alternative splicing into and out of transposons can be highly penetrant

Chimeric transcripts could be inconsequential to a cell if they only constitute a small percentage of the overall transcript repertoire of a given gene. We therefore quantified the percentage of mRNAs produced from a transposon harboring gene that include transposon sequence. To do this we analyzed sites where transposons were spliced into an exon-intron junction of a gene. For each gene we counted the number of reads spanning transposon-exon boundaries, and the number of reads that spanned the exon immediately up- and downstream of the transposon insertion (Figure 6a). This analysis showed that for some genes, all derived mRNAs contained transposon sequences. For example, all spliced copies of the isoform B of the Allatostatin A receptor 1 (*AstA-R1*) contained a section of ROO, and every transcript of *Piezo-like* (*Pzl*), a gene encoding a predicted mechanosensitive ion channel, ended in the I-element instead of exon 1 (Hu et al., 2019; Larsen et al., 2001). All mRNAs retrieved from *Mec2* contained TRANSPAC as the most 5′ sequence, suggesting that transcripts might initiate within the TRANSPAC transposon and spliced to the SD of exon 2 of *Mec2*. On average, transposons contributed to 14% of transcripts derived from a specific gene (Figure 6b, Supplemental Table 7). Insertions in genes on the X chromosome, resulted in a higher average percentage of chimeric transcripts (27% vs. 13% for the rest of the genome) and a larger number of cases where 75% or more of transcripts were chimeric. Insertions on the X chromosome are hemizygous in male flies and our samples were generated by pooling an equal number of male and female flies. The increased chimera rate on the X might therefore reflect the smaller number of different X chromosomes when compared to the rest of the genome. For example, we found that the X-linked *cacophony* (*cac*) gene, which encodes a voltage-gated calcium channel, harbored a sense-orientation BLASTOPIA (Smith et al., 1996). This transposon insertion resulted in 54.7% of *cac* transcripts being truncated in αβCherry fly samples, potentially missing the last 8-11 coding exons, suggesting that many flies in this strain are likely mutant for the *cac* gene (Supplemental figure 7). Another interesting example on the X chromosome of αβCherry flies is *Beadex* (*Bx*) which encodes a long-term memory relevant LIM-type transcription factor (Hirano et al., 2016). A sense NOMAD insertion gives rise to at least two new *Bx* transcript isoforms (Supplemental figure 8), which constitute 9.8% of all *Bx* transcripts.

**Figure 6.**
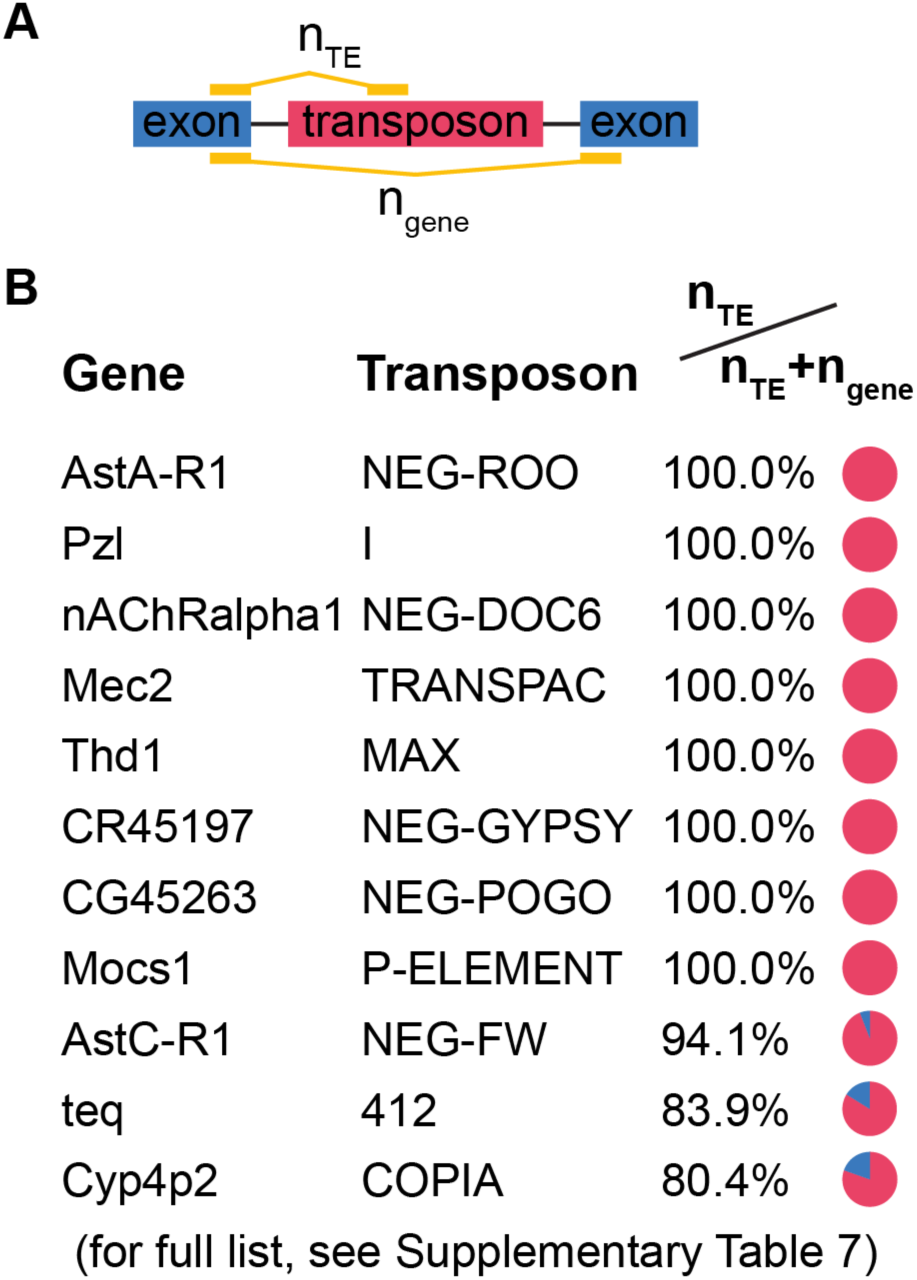
High penetrance of transposon-containing splice isoforms. **A** Schematic showing method used to calculate the frequency of transposon chimera produced from a given gene. The number of reads that span the exon-transposon junction were divided by the total number of reads that map beyond the exon-intron junction (exon-transposon reads plus reads that partially map to the exon immediately downstream of the transposon insertion. **B** Examples of chimeric transposon-gene transcripts detected in fly heads. Many genes are almost exclusively expressed as chimeric transcripts with transposon sequence. See Supplemental Table 7 for complete list.

### Splicing into transposons is common and varies between strains

The transposon complement is highly variable between fly strains. We therefore tested whether other fly strains express chimeric gene:transposon mRNAs by analyzing three previously published mRNA sequencing data sets (Croset et al., 2018; Daines et al., 2011; MacKay et al., 2012). Although these prior studies generated shorter paired-end RNAseq reads, than those collected here, we were still able to find chimeric mRNAs in all three data sets (Supplemental Table 8). Some transposon:gene chimera were conserved across all strains, whilst others appeared to be strain-specific. 318 of the 887 chimera identified in our αβCherry flies were present in at least one of the three other data sets, whereas 120 of those occurred in at least three of the four strains. Chimera that were not detected in data from other strains could either reflect genomic heterogeneity between the tested fly strains, or the absence of evidence could result from lower sequencing coverage. Nevertheless, these results demonstrate the prevalence of cellular mRNAs containing transposon sequence.

## Discussion

Somatic transposition in neurons has been proposed to contribute to age-dependent neuronal decline in wildtype and disease models of *Drosophila* and (Li, e al., 2013; Guo et al., 2018; Sun et al., 2018). Although the frequency of neural transposition is debated, expression is a prerequisite for movement. Therefore, transposons can only mobilize in cells that express full-length elements, or transposon mRNAs that encode enzymes permitting other elements to move in *trans*. It is therefore important to understand how transposon expression is controlled in the brain. Here, we combined single-cell expression data from the *Drosophila* midbrain with high-coverage gDNA sequence data of the same fly strain and find that the majority, if not all, expressed transposon sequences are parts of chimeric mRNAs with cellular genes.

Transposons residing in introns have previously been shown to contribute novel exons to several genes (Nekrutenko and Li, 2001). In addition long non-coding (lnc) RNAs frequently contain transposon sequences (Kapusta et al., 2013). We found transposon exonization to be highly prevalent in the *Drosophila* brain. Transposons within the coding regions of genes and those in introns dramatically increase the transcript repertoire by introducing new splice variants. At this stage it is difficult to definitively determine the whole-genome functional consequences of splicing into transposons because we most often only retrieve the sequence across the splice junctions. Furthermore, although each transposon has a known consensus sequence individual copies are highly polymorphic. Nevertheless, our sequencing shows that transposon exonization often truncates and/or changes the amino acid sequence of the encoded gene products, potentially changing protein structure and function. We did also identify several examples where the inclusion of transposon sequence conserved the reading frame of the host gene, likely generating a novel chimeric protein. Amongst the 148 transposon harboring genes identified in this study, there are several that we have described in detail for which the observed locus disruption and altered expression would be expected to have significant consequences for neural function. Flies harboring HOBO in *Sh* and BLASTOPIA in *cac* might exhibit altered voltage-gated currents, whereas those with ROO in *AstA-R1* will respond differently to the modulatory Allatostatin A neuropeptide (Larsen et al., 2001; Smith et al., 1996). We also described insertions of 412 in *teq* and NOMAD in *Bx*, two genes which have been implicated in long-term memory formation (Didelot et al., 2006; Hirano et al., 2016).

Some transposons provide more splice sites than others. Our analysis of the abundant ROO retrotransposon (found in sense orientation inside 48 genes, and in antisense orientation inside 59 genes in our strain) provides a good example, and the insertion in *mtd* exemplifies the extended variance transposons can introduce to cellular transcription units. Interestingly, antisense insertions of ROO provide more potential SA and SD sites than sense orientation ROO. Insertions of ROO that are sense to the expression of the host gene were found to be mainly part of pre-mRNA, from which functional reverse transcriptase could potentially be translated. In contrast, antisense ROO introduces over 10 times more cryptic splice sites (compare Figure 5b and c).

mRNA molecules that contain transposon sequences might be susceptible to short-nucleotide mediated gene-silencing (Malone and Hannon, 2009), as recently reported for LINE-2 containing mRNAs and LINE-2 derived microRNAs in humans (Petri et al., 2019). This would provide a challenge for the cells expressing transposon-targeting piRNAs in differentiating between chimeric transcripts and full-length transposon sequences. It is worth noting that transposon-directed piRNAs have been identified in the fly brain (Li et al., 2009). It is therefore conceivable that transposon sequence permits cellular mRNAs to be selectively regulated. In addition, transposon sequence might confer the capacity to be specifically trafficked within the cell, and even between cells (Ashley et al., 2018; Pastuzyn et al., 2018).

The process of transposable elements acquiring new cellular functions that benefit the host cell is called transposon exaptation (Gould and Vrba, 2013). Several examples of exaptation events during the evolution of eukaryotes have been reported. How these functional transitions occur is, however, not fully understood (Joly-Lopez and Bureau, 2018). Stress-induced transposon mobilization has been observed in many species, and it has been hypothesized that these mobilization events trigger the formation of new transposon-gene chimera. Our results reveal a new mechanism by which transposons participate in the generation of new gene variants. We show that transposons do not need to mobilize in order to form chimeric mRNA molecules. Instead, the cellular gene splicing machinery frequently uses intronic transposon insertions to increase transcriptional diversity.

Transposon sequences in the gDNA can deliver enhancer elements to cellular genes and thereby influence their expression. However, our results do not support the idea that transposons provide enhancer sequences contributing to neural expression of neighboring genes. If this was the case, we would expect to have found that cellular genes that harbor the same type of transposon are more likely to be expressed in the same cells. Our data instead show that transposon expression is dictated by the genes they are inserted within. This is further supported by the fact that many genes that share transposon insertions with neurally expressed genes were not expressed in the brain.

Our studies also introduce important practical concerns for the analysis of transposon expression in somatic tissue and disease models. Several previous studies have used Quantitative PCR (qPCR) to measure levels of transposon expression in mutant flies and disease models. Since qPCR probes are usually not designed to cross transposon-gene breakpoints, they cannot distinguish autonomous transposon expression from chimeric mRNA (Guo et al., 2018; Li et al., 2013; Sun et al., 2018). Similarly, standard RNA sequencing protocols rely on the alignment of cDNA fragments to the reference genome and would not identify chimeric reads as non-autonomous transposon expression (e.g. De Cecco et al., 2013). Baseline and changing cell-specific expression of host genes that form chimeric transcripts with transposons can therefore be misinterpreted as cell-restricted autonomous transposon expression and mobilization.

Our data also constrain somatic transposition. If all transposon expression depends on the neighboring host gene, only cells expressing that host gene can be susceptible to transposition of that element. This would mean that GYPSY could only be active in glia, if the fly strains being studied harbor a copy of GYPSY in a glial-expressed gene (Krug et al., 2017). Interestingly, although we found sense GYPSY sequences in mRNAs for 14 different genes, we did not detect glial GYPSY expression in our αβCherry flies.

Analysis of RNA-seq data from other fly strains suggests that more than half of the chimeric transposon transcripts we identified in αβCherry flies are unique to this strain. This finding alone demonstrates the incredible heterogeneity of the transposon complement between strains. In addition, our prior genome sequencing revealed large differences between individual αβCherry flies (Treiber and Waddell, 2017). Given the broad range of target genes that we have identified to form chimeric mRNA with transposable elements, it will be important to understand the functional consequences of fixed and variable transposons inside genes for the host organism. Nevertheless, it seems highly likely that polymorphism of transposon load and the distribution of transposons across the host genome could contribute towards heterogeneity of neural function, and neurological pathology, between individual animals.

## Methods

### Fly strains

All experiments were performed on αβCherry flies, which were generated by crossing MB008b females (Aso et al., 2014) with w-; +; UAS-mCherry males. Flies were raised on standard molasses food at 25°C, 40-50% humidity and 12 h:12 h light-dark cycles.

### Long-read mRNA sequencing

For RNA extraction, groups of ∼50 flies were frozen in liquid nitrogen and vortexed for 6 × 30 s to separate heads from abdomens. Heads were isolated using a sieve. To avoid gDNA contamination, mRNA was purified with a combination of protocols. Samples were first processed with a column-based kit (RNeasy Mini kit, Qiagen, UK), including the optional on-column DNAseI digestion. Next, mRNA was extracted from total RNA using oligo-dT magnetic beads (NEB, Ipswich, MA) and the mRNA was purified again using the RNA columns. Finally, sequencing libraries were generated using oligo-dT magnetic beads from a strand-specific mRNA library preparation kit (TruSeq, Illumina, San Diego, CA), with 17 cycles of PCR amplification. Fragmentation was optimized to obtain ∼350nt long fragments. Whole-genome sequencing was performed on a HiSeq 2500, with 250nt paired-end reads. We mapped the long reads using MRTemp, as previously described (Treiber and Waddell, 2017).

### Single-cell read alignments

The *Drosophila melanogaster* reference genome release 6.25 was used for all sequence alignments. Transposon reference sequences were taken from Repbase (Jurka, 2000; Kaminker et al., 2002). Repetitive sequences in the *Drosophila* reference genome were masked using Repeatmasker (Smit et al.). Single-cell sequencing data was processed with a custom-built data processing pipeline, which is available on GitHub. The masked reference genome, as well as a gene reference file (refFlat) with all genes and each unique reference transposon sequence is provided as supplemental data.

### Single-cell data analysis

Digital Gene Expression (DGE) matrices were generated as previously described (Butler et al., 2018). Rscripts can be downloaded as supplemental file 1. In summary, DGE’s were filtered (≥ 800 UMIs, ≥ 400 features) and 8 replicates were merged. Gene and transposon expressions levels were normalized separately. Marker genes were taken from Croset et al. (2018).

### Co-expression analysis

Co-expression was quantified by calculating the Co-expression Disequilibrium (CD, see main text). The R-code snippet is available through GitHub. For the statistical analysis, a non-parametric test was performed. CD values between every gene- and transposon combination were ranked within each biological replicate, p-values were calculated using the student t-test and corrected for multiple comparisons using the Benjamini-Hochberg correction. In addition, CD values were calculated for every tested transposon with a set of 10 randomly assigned genes (or transposons).

### Mapping germline transposon insertions in the fly

Germline transposon insertions were mapped with single-nucleotide resolution using previously published gDNA data from the αβCherry fly strain and MRTemp (Treiber and Waddell, 2017). A new, purpose-built, multi-functional sequence analysis pipeline called TEchim was developed. TEchim has 5 key functions: 1. generation of support files, including a masked reference genome and endogenous intron-exon junctions. (input files: reference genome, list of genes, list of transposon sequences). 2. alignment of un-stranded genomic DNA sequence data of multiple sequencing lanes and multiple biological replicates, detection of chimeric sequence fragments with single-nucleotide resolution, and the generation of summary output tables. 3. alignment of stranded cDNA data, detection of chimeric fragments, quantification of reads 4. generation of matching immobile genetic elements (IGE, see main text), analysis of these IGEs. These data are then used to determine sample-specific detection thresholds. 5. Quantification of LTR-gene and LTR-transposon reads (see Figure 4). Detailed descriptions and manuals are available on Github.

### Splice acceptor (SA) and donor (SD) motif analysis

SA and SD motifs in the *Drosophila melanogaster* reference genome were generated by randomly selecting 500 known SA and SD sites from exons and screening for motifs in these sequences using the MEME suite (Bailey and Elkan, 1994) (Supplemental figure 3). These motifs were then searched in sequence sections across transposon-gene breakpoints using FIMO (Grant et al., 2011).

### Data from previously published studies

Raw single-cell sequencing reads from Croset et al. (2018) was obtained from the NCBI Short Read Archive (SRA https://www.ncbi.nlm.nih.gov/sra) with the accession number: PRJNA428955. Genomic DNA data from Treiber and Waddell (2017) was obtained from the Dryad Digital Repository (https://doi.org/10.5061/dryad.fd930).

## Data Access

All processed data is presented in Supplemental Tables 1-8. All raw sequencing data generated in this study have been submitted to the NCBI SRA with the BioProject ID PRJNA588978. Custom-built software packages can be accessed via GitHub (https://github.com/charliefornia/TEchim and https://github.com/charliefornia/scHardyWeinberg)

## Acknowledgements

We are grateful to other members of the Waddell group for discussion. CT was supported by a Wellcome Trust DPhil studentship. SW is funded by a Wellcome Principal Researcher Fellowship (200846/Z/16/Z), Gatsby Charitable Foundation (GAT3237), ERC Advanced Grant (789274) and Bettencourt–Schueller Foundations.

## Author Contributions

C.D.T. and S.W. conceived the project and wrote the manuscript. C.D.T. performed and analyzed all experiments.

## Supplemental figures

**Supplemental figure 1.**
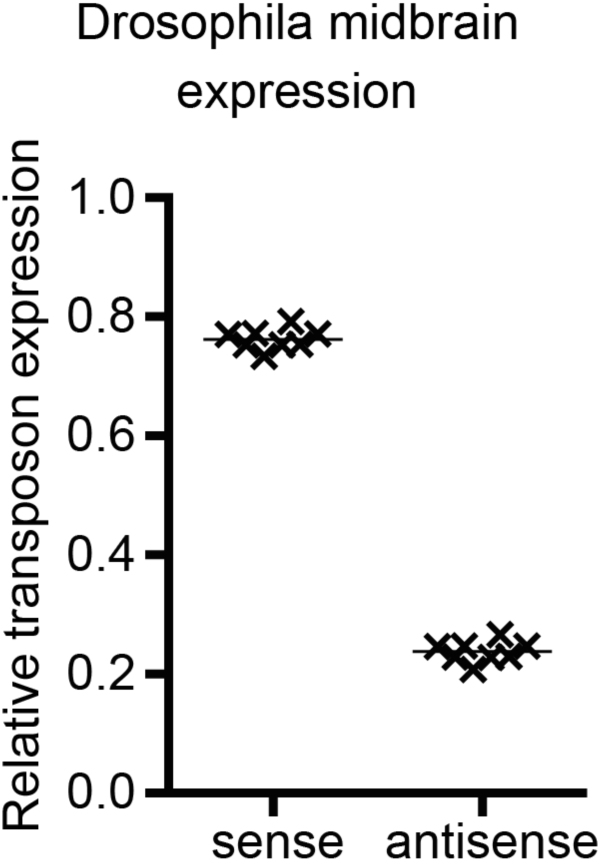
Sense strand transposon transcripts are twice as abundant as antisense strand transcripts in the *Drosophila* midbrain. Graph showing mean expression levels across the entire midbrain of all sense and antisense transposon sequences. Each data point (cross) represents one biological replicate.

**Supplemental figure 2.**
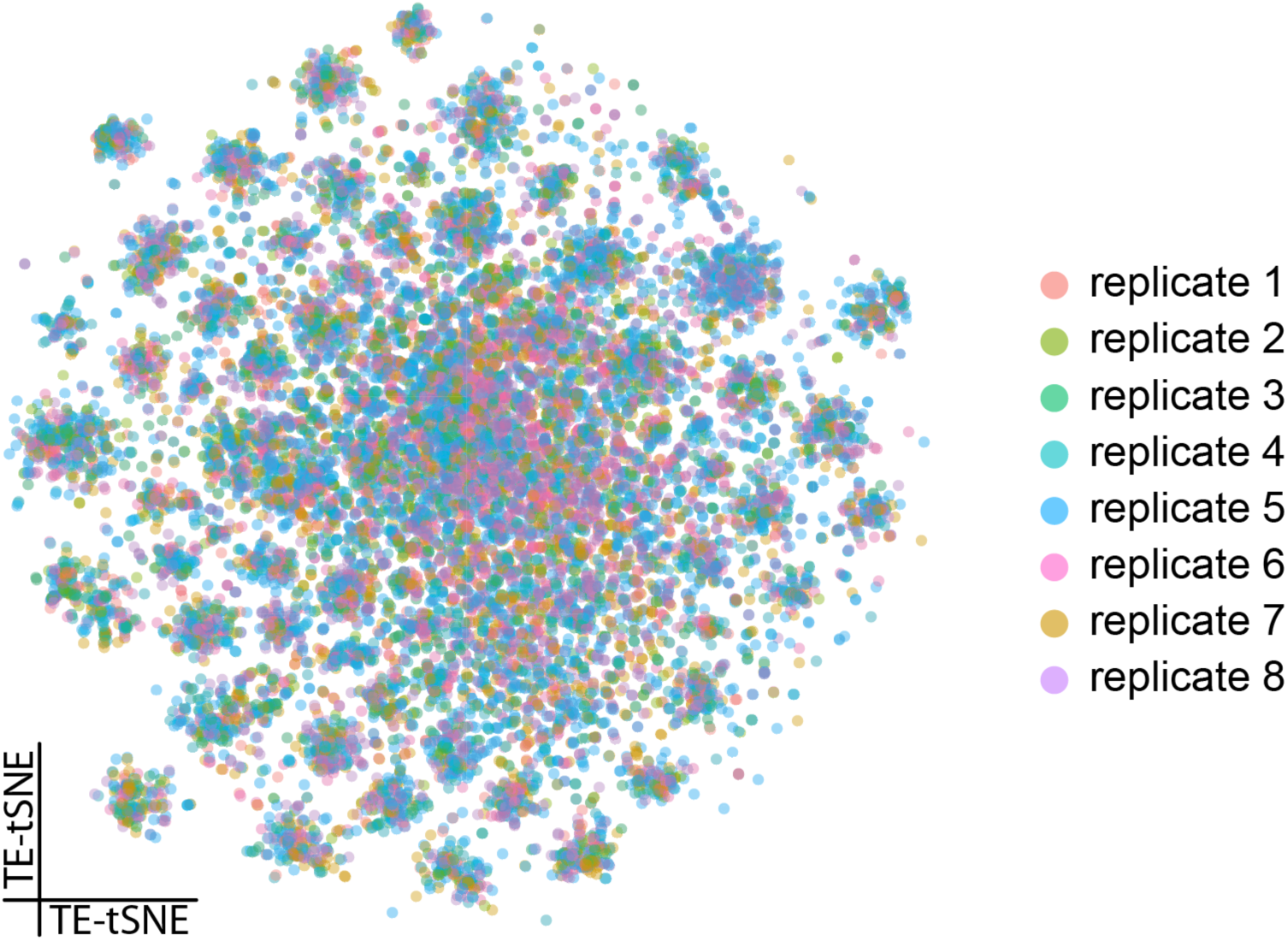
Transposon expression patterns are stereotyped across biological replicates. tSNE based on transposon expression levels showing all 8 biological replicates. Each replicate contributes cells to each cluster.

**Supplemental figure 3.**
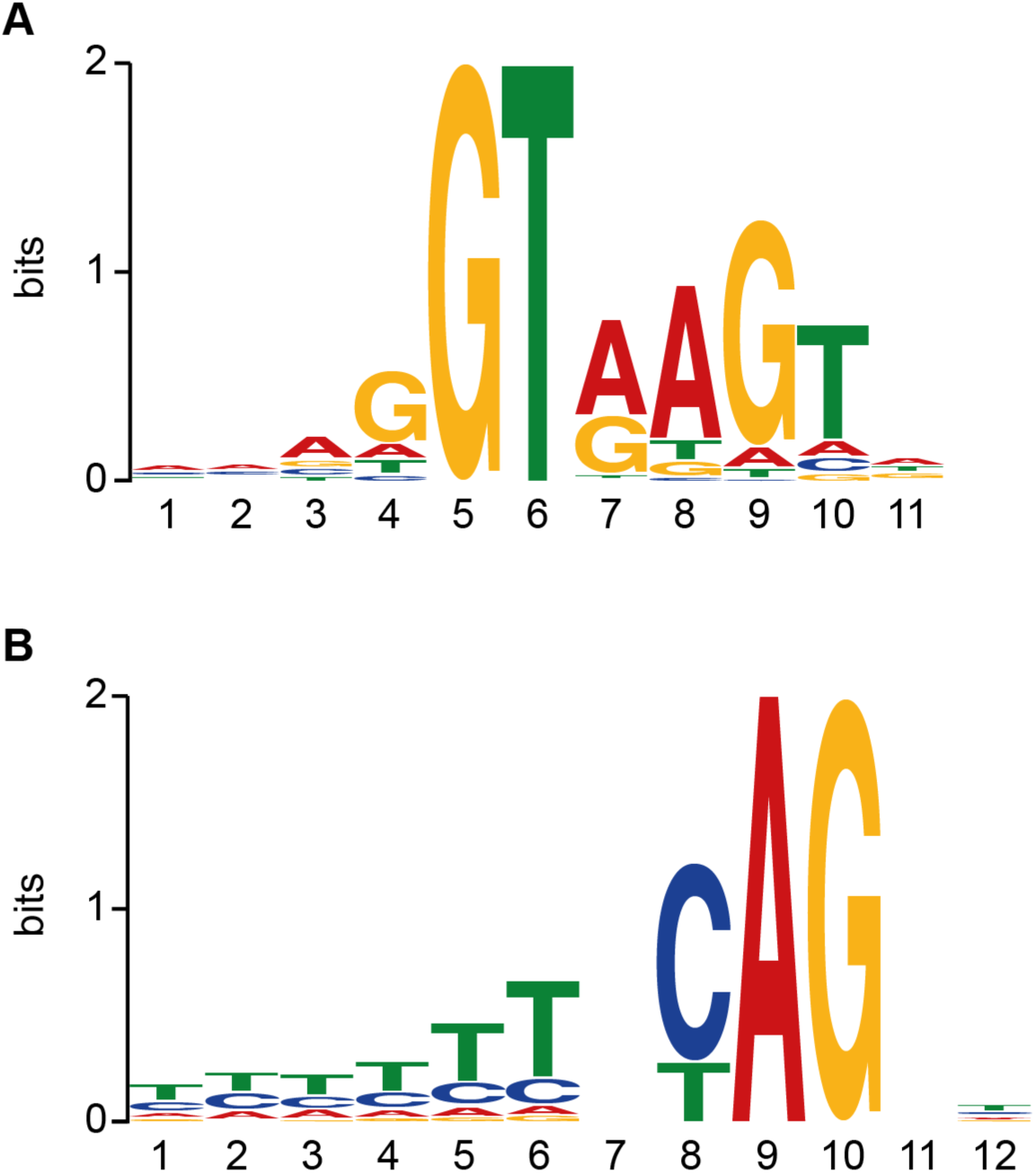
Splice acceptor and donor motifs. **A** Splice acceptor motif, taken from 500 randomly chosen exon-intron junctions of *Drosophila* genes. **B** splice donor motif.

**Supplemental figure 4.**
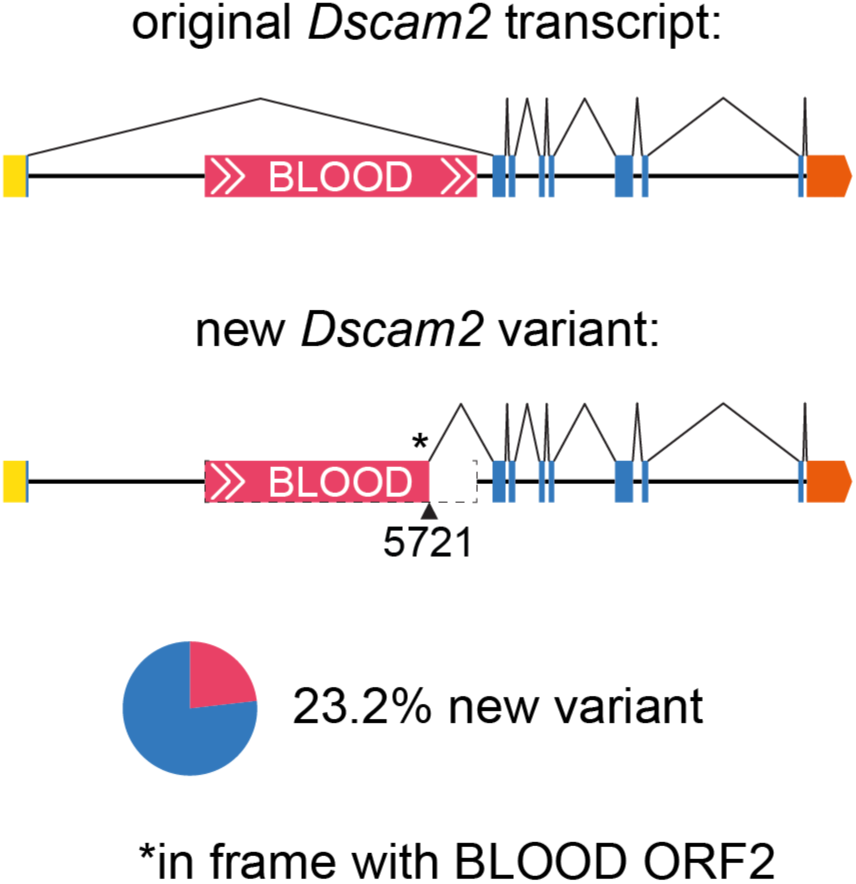
Sense BLOOD insertion in *Dscam2*. Schematics of DScam2 mRNAs produced from locus containing BLOOD. Top shows the nascent transcript spliced around the intronic full-length sense BLOOD insertion. Bottom illustrates a new mRNA splice isoform, which reads through in frame from the ORF2 sequence of BLOOD into exon 2 of *DScam2*. The breakpoint in BLOOD is a consensus SD motif. 23.2% of all *Dscam2* transcripts in fly heads are chimeric with BLOOD.

**Supplemental figure 5.**
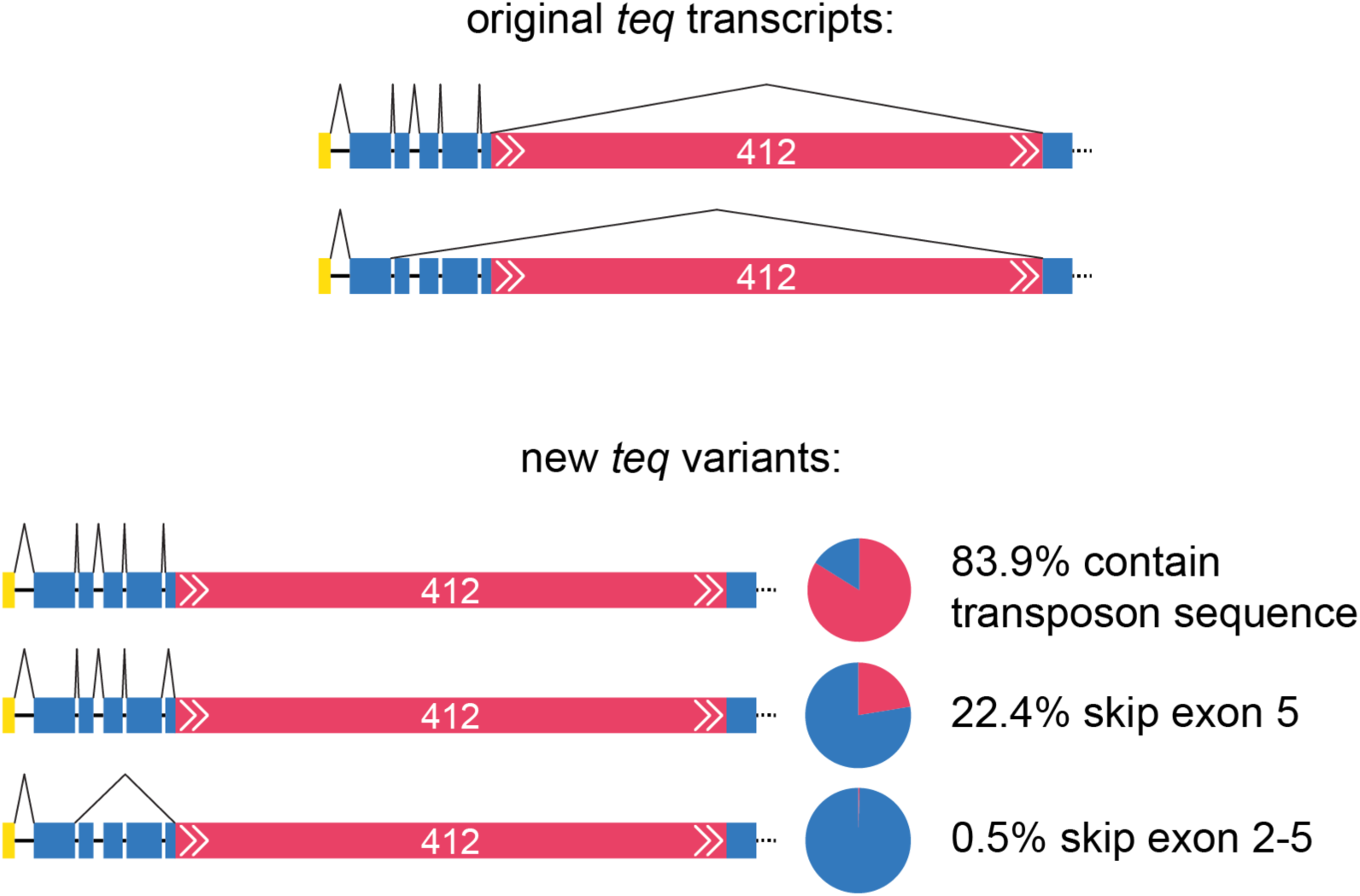
Sense 412 insertion in *tequila*. Schematics of *teq* mRNAs produced from locus containing full-length sense orientation 412 insertion. Top, original transcripts of *teq* splicing around 412. Bottom, and new *teq* splice isoforms that include 412 sequence. 83.9% of *teq* transcripts contain 412 transposon sequence. In addition, 22.4 % of 412 containing mRNAs skip exon 5 of *teq*. In 0.5% of cases exons 2-5 are skipped.

**Supplemental figure 6.**
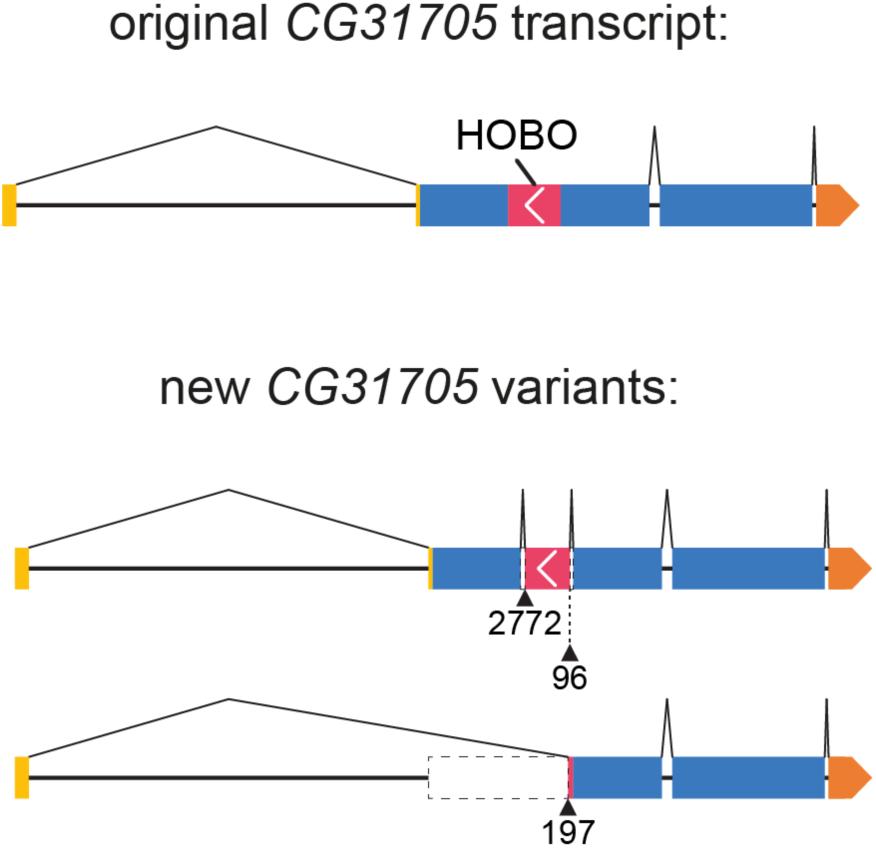
Antisense HOBO insertion in *CG31705*. Schematic of *CG31705* mRNAs produced from locus containing exonic antisense HOBO insertion. Transcripts containing unspliced HOBO and two additional new splice isoforms that are generated by alternative splicing into HOBO are shown.

**Supplemental figure 7.**
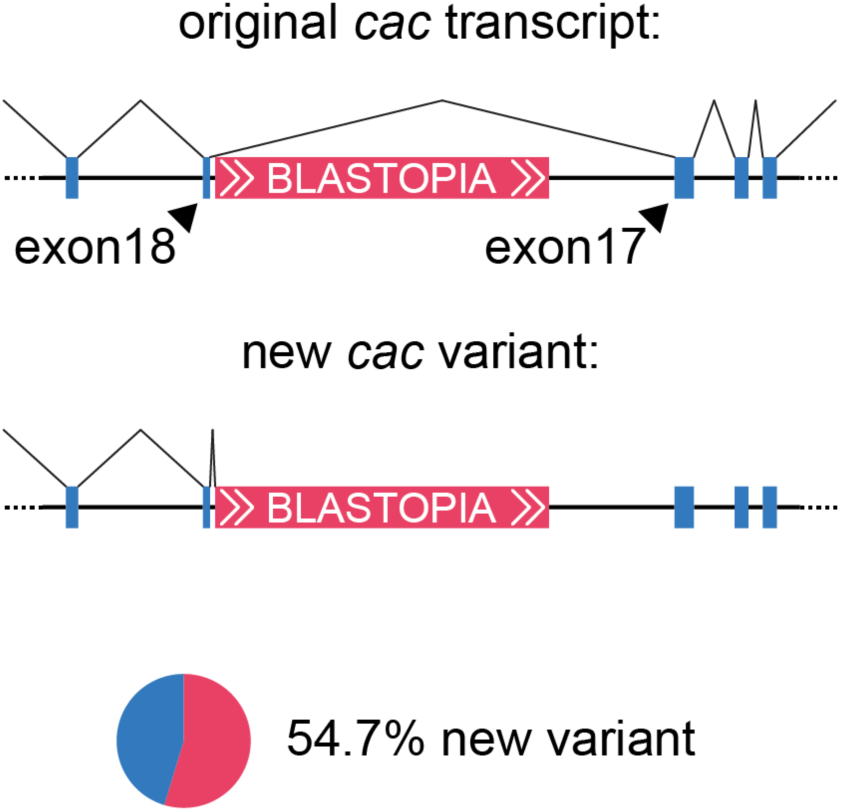
Sense BLASTOPIA insertion in *cacophony*. Schematic *cac* transcripts produced from locus containing a full-length intronic sense BLASTOPIA insertion (the orientation is 5′ (left) to 3′ (right)). A regular cac transcript is produced by splicing around the BLASTOPIA insertion and a new truncated *cac* isoform results from splicing into BLAsTOPIA. The *cac* gene is on the X chromosome, and 54.7% of *cac* transcripts in the fly head are truncated by splicing into BLASTOPIA.

**Supplemental figure 8.**
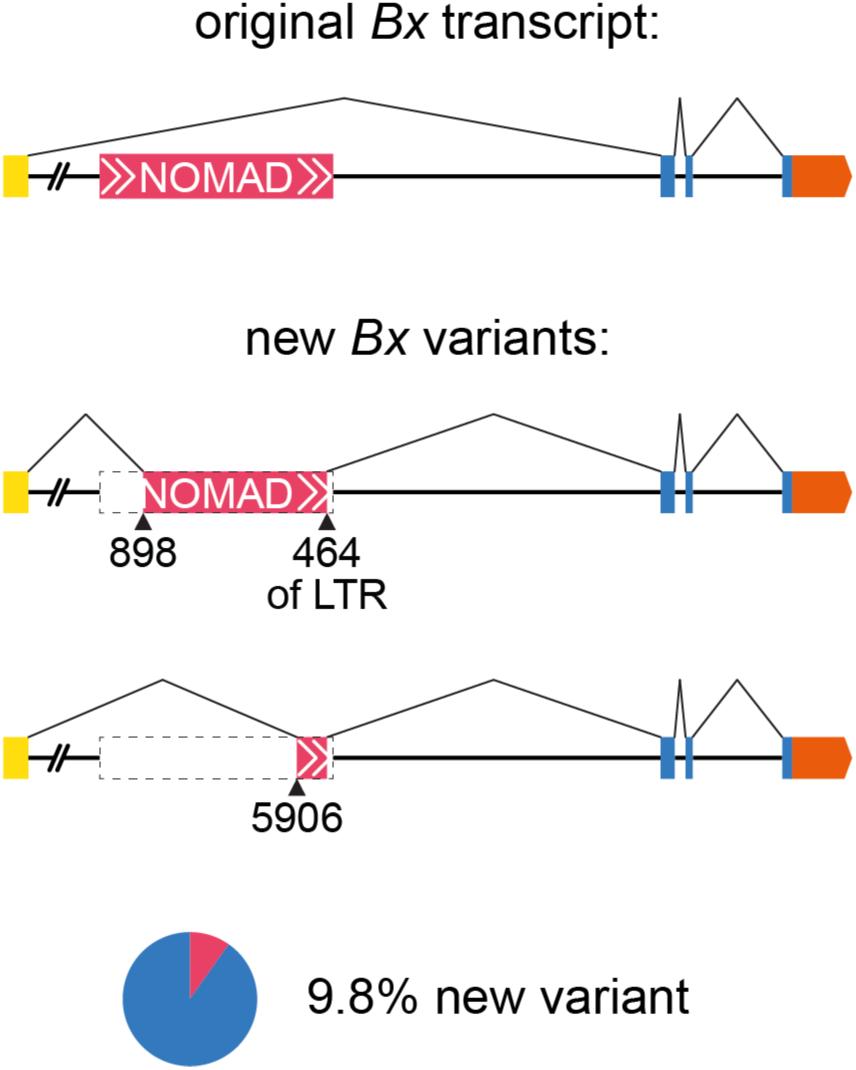
Sense NOMAD insertion in *Bx*. Schematic showing mRNAs produced from locus containing sense intronic NOMAD insertion. Original transcript of *Bx* is generated by splicing around NOMAD. New splice isoforms contain fragments of NOMAD. 9.8% of transcripts that start with the *Bx* 5′UTR are spliced into NOMAD.

